# DEVIATIONS IN WHOLE BODY ANGULAR MOMENTUM ARE LARGELY CORRECTED BEFORE FOOT PLACEMENT

**DOI:** 10.64898/2025.12.03.692080

**Authors:** Simone Berkelmans, Sjoerd M. Bruijn, Maarten Afschrift

**Author notes:** **Correspondence address:** Simone, Berkelmans, Department of Human Movement Sciences, Faculty of Behavioural and Movement Sciences, Vrije Universiteit Amsterdam, Van der Boechorststraat 9, 1081 BT Amsterdam, The Netherlands.

## Abstract

This study examined how mediolateral foot placement is controlled following mechanical perturbations that affected either whole-body linear or angular momentum. Predictive foot placement models based on center of mass state alone were compared with models that additionally included whole-body angular momentum to determine whether whole-body angular momentum contributes to foot placement control beyond linear momentum. Ten healthy adults walked on a treadmill at 2 km/h and 5 km/h while being exposed to two perturbation types: (1) a pull to the pelvis that primarily altered linear momentum (translation perturbation) and (2) simultaneous pulls to the pelvis and shoulder in opposite directions that primarily altered angular momentum (rotation perturbation). Perturbations were applied at heel strike, with a magnitude of ∼120 N and a duration of 300 ms. Whole-body kinematics were recorded using 3D motion capture and processed in OpenSim to compute linear and angular momentum. Translation perturbations caused large deviations in whole-body linear momentum with minimal changes in whole-body angular momentum, whereas rotation perturbations induced strong whole-body angular momentum deviations with smaller changes in linear momentum. Including whole-body angular momentum minimally improved foot placement predictions during early swing after rotation perturbations. These findings indicate that mediolateral foot placement is primarily governed by linear momentum dynamics.

## Introduction

In human walking, the control of linear momentum, particularly that of the center of mass (CoM) motion with respect to the stance foot, has been extensively studied. Humans regulate center of mass motion by modulating the ground reaction force, which, aside from gravity, is the only external force acting on the body (Hof, 2007; Reimann et al., 2018; Winter, 1995). This modulation occurs both through stance-phase strategies, such as ankle and hip torque, and through foot placement (Bauby & Kuo, 2000; Bruijn & Van Dieën, 2018; MacKinnon & Winter, 1993; Reimann et al., 2019; Townsend, 1985). Empirical studies have shown that foot placement location is closely related to the CoM’s position and velocity at mid-stance (Van Den Bogaart et al., 2020; Vlutters et al., 2016; Wang & Srinivasan, 2014). During steady-state gait, muscle activity and the resulting ground reaction force vectors are finely tuned to stabilize the CoM trajectory and maintain balance (Hof & Duysens, 2018; Reimann et al., 2019). This relationship between CoM dynamics and foot placement, has been formalized through the concept of the extrapolated Center of Mass (XCoM), which combines the instantaneous CoM position and velocity with the dynamics of an inverted pendulum model (Bruijn et al., 2013; Hof, 2007; Hof et al., 2005). The XCoM framework has successfully estimated step width and foot placement adjustments, providing insight into how humans anticipate and correct for potential instability in the sagittal and frontal plane (Bruijn & Van Dieën, 2018; O’Connor & Kuo, 2009; Van Mierlo et al., 2022).

In addition to linear momentum, control of angular momentum about the CoM (throughout the manuscript defined with respect to the whole-body center of mass) is a requirement for stable walking (Herr & Popovic, 2008; Van Dieën et al., 2025). While linear momentum provides information about the translation of the body, it does not capture the rotational dynamics of the body. Both linear and angular momentum are determined by external forces, primarily gravity and ground reaction forces, with the latter modulated through stance-phase ankle torques and foot placement (Reimann et al., 2019; Van Den Bogaart et al., 2020). Since gravity is constant, humans achieve this control by adjusting the magnitude, direction, and point of application of ground reaction forces to maintain a stable, periodic gait (Herr & Popovic, 2008; Hof, 2007).

Although foot placement is often viewed as a mechanism to modulate the ground reaction force for controlling the motion of the CoM (Bruijn & Van Dieën, 2018; Van Den Bogaart et al., 2020), Leestma et al. (2023) found that step placement adjustments is also correlated with variations in integrated whole-body angular momentum, particularly in the frontal plane. Complementing this, Van Dieën et al. (2025) proposed that humans use simultaneous feedback control of linear and angular momentum during walking, implying that foot placement is also used to control angular momentum. These findings suggest that foot placement is shaped by the combined need to regulate both linear and angular momentum, rather than linear momentum alone.

Experimental evidence indicates that, under certain conditions, humans prioritize the control of angular over linear momentum. For instance, Pijnappels et al. (2004) showed that following a trip, humans often first control trunk rotation and restore whole-body angular momentum at the cost of deviations in center of mass velocity, highlighting the prioritization of controlling angular momentum. Similarly, Van Mierlo et al. (2022) demonstrated that responses to angular momentum perturbations occur substantially faster than responses to linear perturbations in the sagittal plane. These findings underscore that in challenging situations angular momentum regulation is not only an integral part of locomotor control but can also dominate stabilizing strategies.

While previous work has established that foot placement is tightly coupled to the CoM state and can be well predicted by models such as the XCoM (Hof, 2007; Vlutters et al., 2016; Wang & Srinivasan, 2014), it remains less clear whether this relationship fully captures how humans control stability when whole-body angular momentum is perturbed. In the present study, we investigated how mediolateral foot placement is controlled following mechanical perturbations that selectively affected either CoM linear momentum (translation perturbations) or whole-body angular momentum (rotation perturbations). By comparing predictive models based on CoM state alone with models that additionally included whole-body angular momentum, we aimed to determine whether whole-body angular momentum contributes to the control of mediolateral foot placement beyond linear CoM dynamics. We hypothesized that incorporating whole-body angular momentum would significantly improve prediction of mediolateral foot placement following rotational perturbations, as these perturbations directly alter the body’s rotational state. In contrast, we expected that models based solely on CoM state would sufficiently explain foot placement following translation perturbations. In addition to these hypothesis-driven analyses, we conducted exploratory analyses to characterize participants’ responses during and immediately following the perturbations to further interpret the control strategies used to maintain balance.

## Methods

### Participants

Fourteen healthy participants (four men; age 23.6 ± 1.9 years; weight 69.6 ± 10.2 kg, height 1.72 ± 0.06 m, mean ± SD) participated in the study. Four participants were excluded due to missing marker data or missing load cell data, resulting in ten participants being included in the analysis. Exclusion criteria were any sports injuries or musculoskeletal conditions that could affect walking patterns or balance, as these could influence the results. The study was approved by the local ethics committee (VCWE-2024-065), and all participants provided written informed consent prior to participation.

### Measurement setup and data collection

Participants walked on a large treadmill (250 cm × 450 cm; Rodby innovation, RL4500E, Sweden; Figure 1). A projector displayed the outline of a standard-size treadmill (75 cm × 200 cm) onto the large treadmill surface, creating a “small treadmill” within the larger belt area. For the present study, only perturbations on the large treadmill were analyzed. Participants wore a full-body safety harness attached to an overhead safety rope to prevent injury in case of a fall and to ensure they would not contact the treadmill surface.

**Figure 1.**
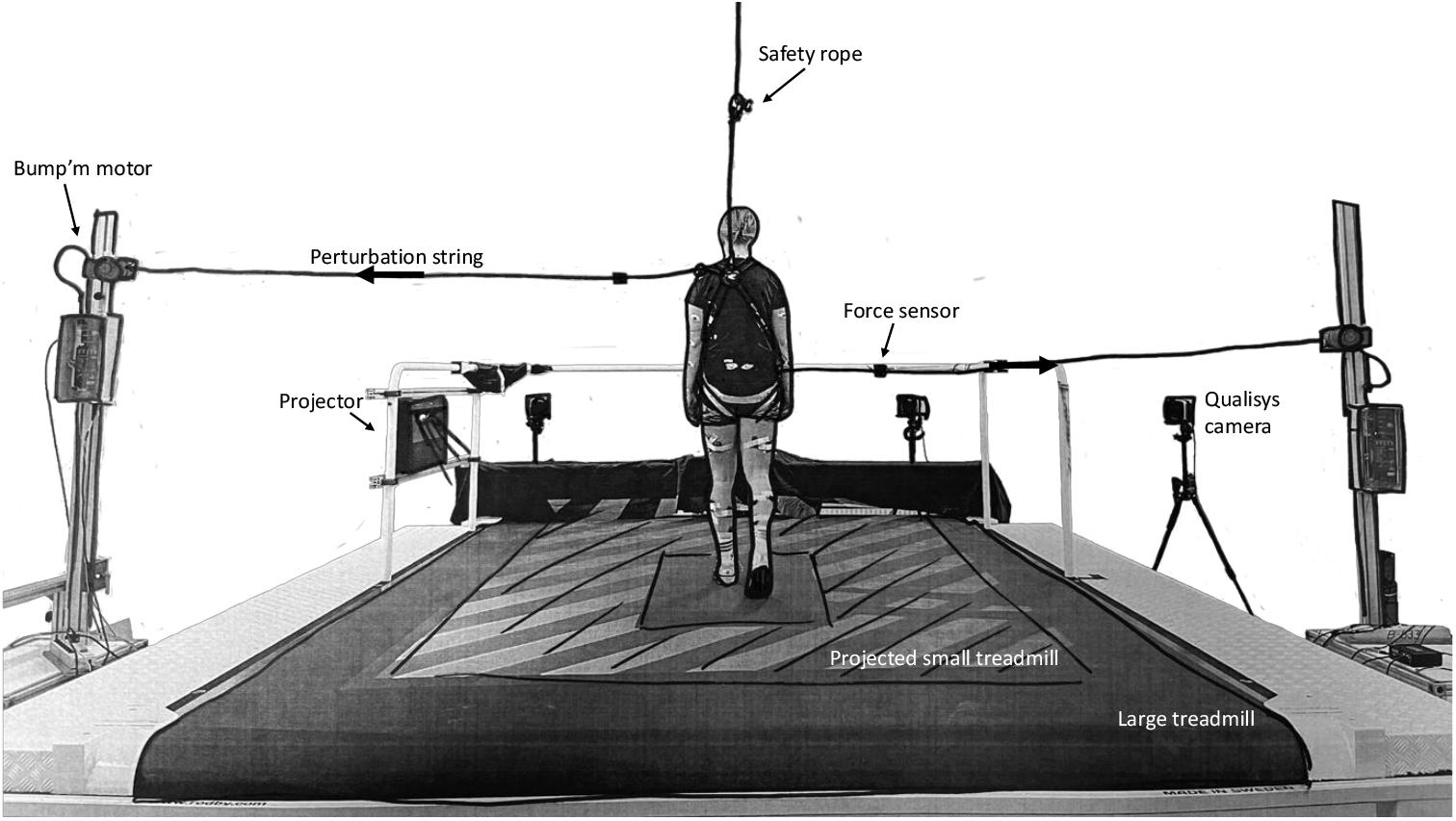
Measurement setup.

Mechanical perturbations were applied using two motors (Bump’m; Tan et al., 2020). The right motor was mounted approximately 75 cm above the treadmill base, approximately aligned with the participant’s pelvis, to deliver right-lateral pulls. The left motor was mounted approximately 150 cm above the treadmill base, approximately aligned with the left shoulder, to deliver left-lateral pulls. Each motor was connected to the harness via perturbation strings attached at the right pelvis and left shoulder, respectively. The motors were positioned approximately 1.5 m horizontally from the participant, resulting in a nearly horizontal rope direction (Figure 1). A force sensor (Avanti Load Cell Adapter, Delsys Inc. Natick, MA, USA; sampling frequency 2000 Hz) was in-line with each perturbation string to record the applied loads. To ensure consistency, we focused on the net moments acting on the CoM. Inspection of these net CoM moments across participants (see Supplementary Figure S15) confirmed that the perturbations produced comparable effects.

Seven Qualisys Argus cameras (Qualisys, Göteborg, Sweden; sampling frequency 100 Hz) captured full-body kinematics. Reflective markers were placed on anatomical landmarks and cluster locations (Supplementary Material, Table S1), including the harness and perturbation strings to track the applied forces. All systems were synchronized using a 5V pulse from the Qualisys sync unit.

### Experimental protocol

All participants walked on the treadmill at two speeds: slow (2 km/h, ∼0.56 m/s) and normal (5 km/h, ∼1.39 m/s). Prior research has shown that deviations in CoM state and deviations in foot placement might be speed dependent, which is particularly relevant for clinical populations such as individuals with spinal cord injury (SCI), who often walk at significantly reduced speeds (Zadravec et al., 2020). The experiment was divided into two blocks (slow and normal walking speed), with all participants starting with the slow walking speed condition. Within each block, participants walked on both the large and the projected small treadmill, with the order of treadmill type randomized across participants. Participants were instructed to respond to perturbations by regaining balance and returning to their original position on the treadmill.

Before starting the first block, participants completed a five-minute familiarization period to adjust to walking on the treadmill as well as to the safety harness and string attachment. During this period, they walked at both walking speeds.

Perturbations were applied either as a force to the pelvis alone or as rotation perturbation involving simultaneous forces to both the pelvis and shoulder (Figure 2A and B). The rotation perturbation generated a moment on the torso without producing a net force (see Figure 2C). Each block included four perturbation conditions: Translation Right, which was a right pelvis pull applied at initial contact of the right leg; Translation Left, a right pelvis pull at initial contact of the left leg; Rotation Right, a combined pull on the right pelvis and left shoulder at initial contact of the right leg; and Rotation Left, the same combined pull applied at initial contact of the left leg. Each perturbation was an impulse lasting 300 ms with a force magnitude of ±120 N, comparable to the strongest perturbations used in Van Mierlo et al. (2022). Each condition was repeated 10 times per block, resulting in a total of 160 perturbations across the entire experiment. Perturbations were delivered at random intervals ranging from 8 to 12 seconds to avoid anticipation.

**Figure 2.**
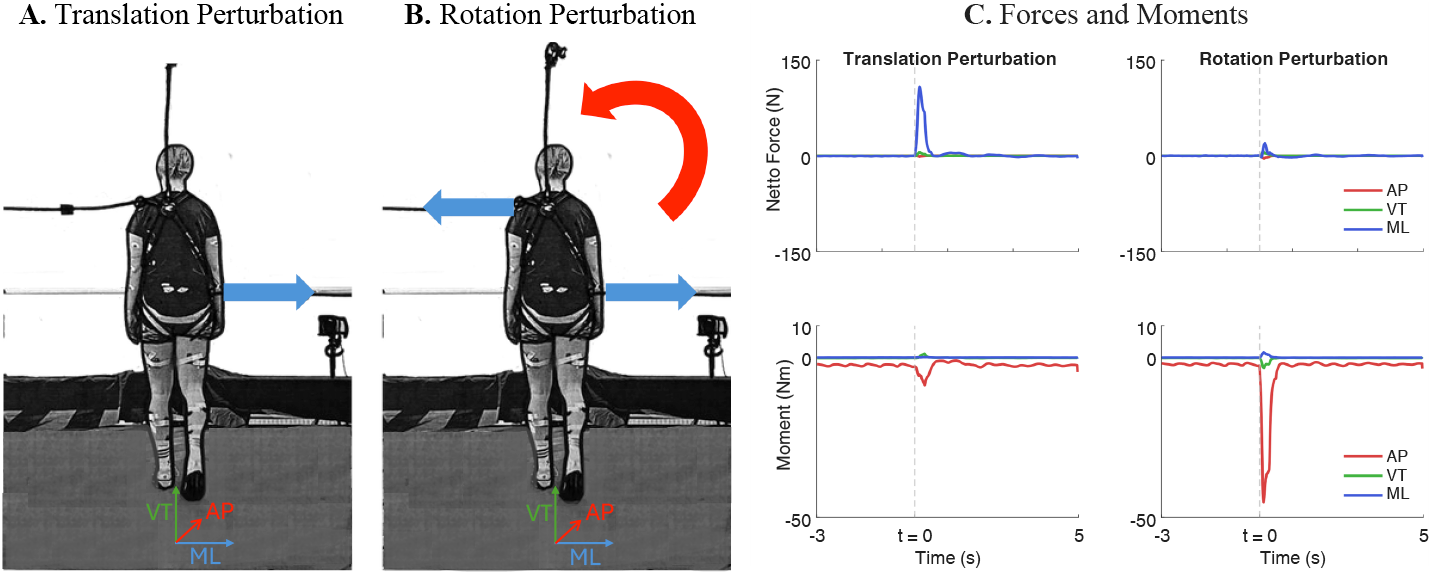
(A) Translation perturbation, applied as a pull (blue arrow) to the pelvis (right); (B) Rotation perturbation, applied as simultaneous pulls (blue arrows) to both the pelvis (right) and shoulder (left), with the resulting moment about the CoM (red arrow); (C) Forces (top panels) and moments about the CoM (bottom panels) during translation (pelvis) and rotation (pelvis–shoulder) perturbations, plotted with perturbation onset at t = 0, including 3 seconds prior and 5 seconds following the perturbation. AP: anterior-posterior, ML: mediolateral, and VT: vertical.

### Data processing

Raw marker data from Qualisys were labeled in QTM-software. The marker trajectories, load cell signals, surface EMG, and perturbation data were preprocessed in Python (Python, version 3.12). For each participant, trials were synchronized across the Qualisys motion capture, load cell recordings, surface EMG, and Bump’m system. All trial data were time-normalized relative to the perturbation onset (*t* = 0). For each trial, three seconds of data prior and five seconds following the perturbation were extracted for analysis.

Gait events were detected from heel and toe marker trajectories (low-pass filtered with a second-order Butterworth, cut off 4 Hz). Heel strikes were identified by locating downward peaks in the vertical velocity of the heel marker and then selecting the first frame after each peak at which the heel vertical velocity returned to approximately zero, corresponding to foot contact. Toe-offs were defined as the first frame after each heel strike at which the anterior-posterior velocity of the toe marker, corrected for treadmill speed, exceeded a predefined threshold. Default thresholds were 0.05 m·s^−1^ for both the heel vertical and toe anterior-posterior velocities unless otherwise specified.

We scaled the generic Gait2392 musculoskeletal model in OpenSim (version 4.5) to each participant based on individual anthropometry and body mass. The scaled model, together with the processed marker trajectories, was used for inverse kinematics to compute joint angles. From these analyses, segment orientations and CoM positions and velocities were derived. Whole-body angular momentum was calculated following Herr & Popovic, (2008):

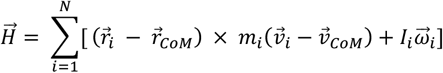

Where *i* represents each body segment of the OpenSim model, *r*_*i*_ and *r*_*CoM*_ are the positions of the *i*^th^ segment CoM and the whole-body CoM, respectively *v*_*i*_ and *v*_*CoM*_ are the velocities of the *i*^th^ segment and the whole-body CoM, *m*_*i*_ is the mass of the *i*^th^ segment, *I*_*i*_ is the segment’s inertia tensor, and *ω*_*i*_ is the angular velocity of the *i*^th^ segment’s CoM (Herr & Popovic, 2008).

### Outcome measures

*Step width, step time*, and *margin of stability* were analyzed for the final unperturbed step (Pre) and the first three steps following the perturbation (Post 1–Post 3). *Step width* was defined as the mediolateral distance between left and right heel markers at heel strike. *Step time* was defined as the temporal interval between heel strikes of alternating feet. Mediolateral *XCoM* was computed according to Hof (2007):

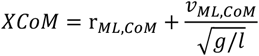

Where *r*_*ML,CoM*_ is the mediolateral CoM position, *v*_*ML,CoM*_ is the mediolateral CoM velocity, *g* = 9.81 *ms*^−2^, and *l* is the effective pendulum length approximated by the mean vertical CoM height (Hof, 2007). The *margins of stability* in the mediolateral direction was then calculated as the distance between the XCoM and the position of the leading foot at toe-off of the trailing foot. A positive margin of stability indicates that the XCoM lies within the base of support.

Mediolateral foot placement was estimated using linear regression models to evaluate if foot placement is related to preceding mediolateral CoM position 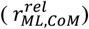 and velocity (*v*_*ML,CoM*_) with respect to the stance foot, and whole-body angular momentum (*WBAM*_*AP*_), corresponding to the rotation around the anterior-posterior axis. The swing phase was time-normalized, and for each time point *i*, a regression model was fitted across all available steps (1,…,total number of steps), using the CoM state (and whole-body angular momentum) at each time point *i* to predict foot placement at heel strike.

Model 1 only included CoM position and velocity as predictors:

1. *Foot Placement* 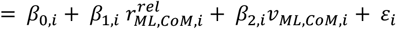 Model 2 additionally included whole-body angular momentum as a predictor:
2. *Foot Placement* 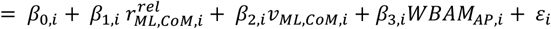

Regression models 1 and 2 were fitted separately for each participant using a least-squares procedure on all steps (approximately 1000 steps per participant) from three seconds before to five seconds after perturbation onset, including both rotation and translation perturbations. For each swing time point, regression coefficients (*β*_0_, *β*_1_, *β*_2_, *β*_3_), relative explained variance (*R*^2^), and step-by-step prediction errors (residuals, *ε*), we obtained. Group-level results were calculated as the average regression coefficients and explained variance across participants. By comparing the model with and without angular moment information, we evaluated how much the additional variance in foot placement is explained when whole-body angular momentum is included alongside CoM kinematics. To quantify differences in model performance across the swing phase, the explained variance (R^2^) of Models 1 and 2 was statistically compared using *SPM1D* (one-dimensional statistical parametric mapping).

### Statistics

*Step time, step width*, and *margin of stability* were analyzed for the final unperturbed step (Pre) and the first three steps following the perturbation (Post 1–Post 3). Separate paired t-tests, or non-parametric equivalent, were performed to compare Pre vs. Post 1, Pre vs. Post 2, and Pre vs. Post 3. This approach was chosen to directly assess how *step time, step width*, and *margin of stability* changed following perturbation exposure, without modeling complex interactions across perturbation types or walking speeds. Walking speed and perturbation type were not included, as the primary interest was in the recovery steps rather than between-perturbation differences. The significance level was set at α = 0.05, and p-values were Bonferroni-corrected for multiple comparisons.

## Results

During translation perturbations, both CoM position and velocity showed substantial deviations from unperturbed steps (see Figure 3), with mediolateral CoM displacements (∼0.2 m) and increases in velocity (up to ∼0.4 m/s). Whole-body angular momentum remained mostly unchanged compared with unperturbed steps, with some slight deviations in the first two post-perturbation steps. These responses were similar between walking speeds. Rotational perturbations caused clear whole-body angular momentum deviations (up to ∼5 kg m^2^/s). CoM position and velocity also deviated from unperturbed trajectories (∼0.05 m), although to a smaller extent than during translation perturbations, especially during the first two post-perturbation steps. Faster walking amplified the whole-body angular momentum response and slightly increased CoM position and velocity deviations relative to slow walking speed.

**Figure 3.**
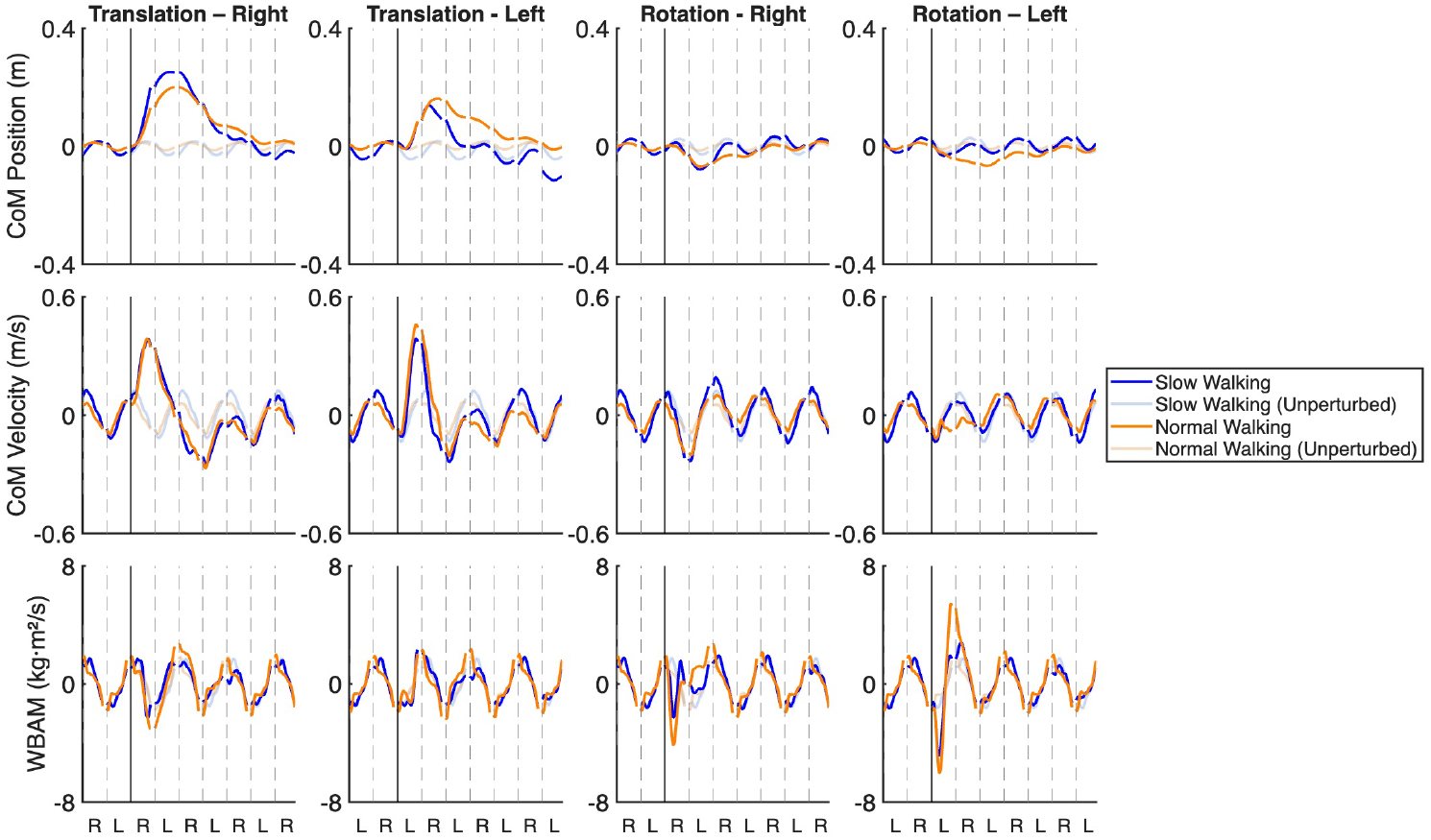
Mediolateral CoM position (top row), CoM velocity (middle row), and whole-body angular momentum (bottom row) across steps for different perturbation types. Columns correspond to perturbation types: Translation Right, Translation Left, Rotation Right, Rotation Left. Blue and orange lines indicate slow and normal walking speed, respectively, with lighter grey shades representing unperturbed steps. Heel strikes are denoted by R (right) and L (left); intervals from R to L indicate right stance, and from L to R indicate left stance. The solid vertical line marks the onset of the perturbation, and dashed vertical lines indicate subsequent heel strikes.

### Step time

Perturbations affected step time (Figure 4A) during both slow and normal walking speed. During slow walking speed, the first post-perturbation step (Post 1) was significantly shorter than the unperturbed step for Translation Right (p < 0.001), Translation Left (p < 0.001), and Rotation Right (p < 0.01), indicating an immediate adjustment in step duration. During normal walking speed, Post 1 step time was significantly reduced for Translation Right, Translation Left, and Rotation Left (p < 0.05), while Rotation Right did not differ from unperturbed (p = 1.000). By Post 2, step time largely returned to baseline. During slow walking, only Translation Left remained different from unperturbed (p < 0.05); no differences were observed during normal walking. By Post 3, step time did not differ from unperturbed for any perturbation type at either walking speed (all p ≥ 0.314).

**Figure 4.**
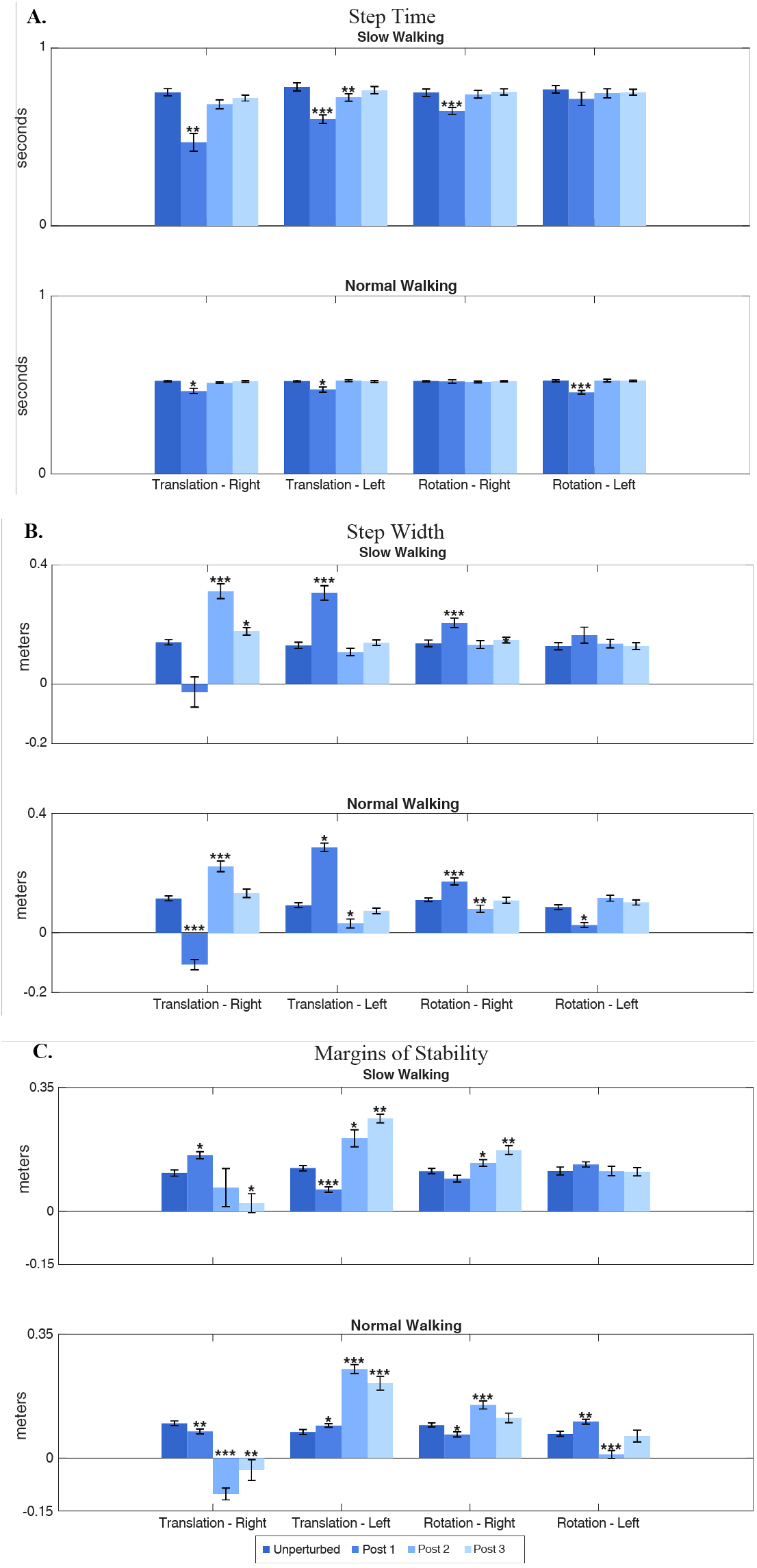
(A) Step time, (B) Step width, and (C) Margins of Stability during slow and normal walking after translation and rotation perturbations. Different colored bars show group mean ± SD for the unperturbed step (Unperturbed) and the first three post-perturbation steps (Post 1–3) for each perturbation type (Translation Right, Translation Left, Rotation Right, and Rotation Left). Step width was defined as the mediolateral distance between left and right heel markers at heel strike. In our analysis, this distance is computed relative to the stance foot. As a result, when a participant crosses one foot in front of or behind the other (a “crossover step”), this relative distance can be negative. * p < 0.05; ** p < 0.01; *** p < 0.001.

### Step width

Perturbations significantly affected step width (Figure 4B) during both slow and normal walking speeds. In general, participants stepped in the direction of the perturbation/applied pull during Post 1 (Post 1, slow walking speed: Translation Left and Rotation Right p < 0.01; normal walking speed: all p < 0.05). Depending on the stance leg at perturbation onset, this resulted in an outward/wider step width (Perturbations: Translation Left, Rotation Right) or a cross over/smaller step width (Perturbations: Translation Right, Rotation Left. For translation perturbations, this directional response persisted into Post 2 at both walking speeds (p < 0.05). In contrast, step width following rotational perturbations returned to baseline by Post 2. By Post 3, step width no longer differed from unperturbed for any perturbation type or walking speed (all p ≥ 0.190).

### Margins of stability

During Post 1, margins of stability (Figure 4C) were significantly reduced relative to the unperturbed step during slow walking for Translation Right and Translation Left (p < 0.05), whereas rotational perturbations did not differ. During normal walking, margins of stability differed significantly from unperturbed for all perturbation types (p < 0.05). The reduction in margins of stability during Post 1 reflects a shift of the margins of stability toward the stance limb, with the direction of change depending on perturbation type and stance limb (outward shift: Translation Left, Rotation Right; inward shift: Translation Right, Rotation Left).

Following translation perturbations, margins of stability remained significantly different from unperturbed in Post 2 for Translation Left during slow walking (p = 0.041) and for both translation perturbations during normal walking (p < 0.001). Differences persisted into Post 3 for both translation directions at both walking speeds (slow: p < 0.05; normal p < 0.01), indicating a prolonged recovery. For rotational perturbations, margins of stability remained altered in Post 2 for Rotation Right during slow walking (p < 0.05) and for both rotation directions during normal walking (p < 0.001). By Post 3, only Rotation Right during slow walking remained different (p < 0.01), whereas all other rotational conditions returned to baseline (p ≥ 0.233), demonstrating a more rapid recovery compared to translation perturbations.

### Foot placement

For rotation perturbations, models including whole-body angular momentum as a predictor (Figure 5A left panel, solid red lines) showed significantly higher (p<0.001) explained variance (R^2^) than models without whole-body angular momentum between ∼10–20% of the swing phase during normal walking speed. In the remainder of the swing phase, R^2^ values were not significantly different between models, indicating that whole-body angular momentum contributes primarily during early swing after rotational perturbation. For translation perturbations, inclusion of whole-body angular momentum (Figure 5A, solid purple lines) did not significantly improve model performance, as R^2^ values were not significantly different from models excluding whole-body angular momentum throughout the swing phase. This pattern was further reflected in the regression coefficients of whole-body angular momentum (β^_3_^, Figure 5A right panel), showing a limited and time-specific influence on foot placement adjustments. This suggests that whole-body angular momentum plays a limited and time-specific role, primarily during early swing after rotational perturbations, and its contribution to foot placement adjustments is largely negligible during slow walking speed and translational perturbations.

**Figure 5.**
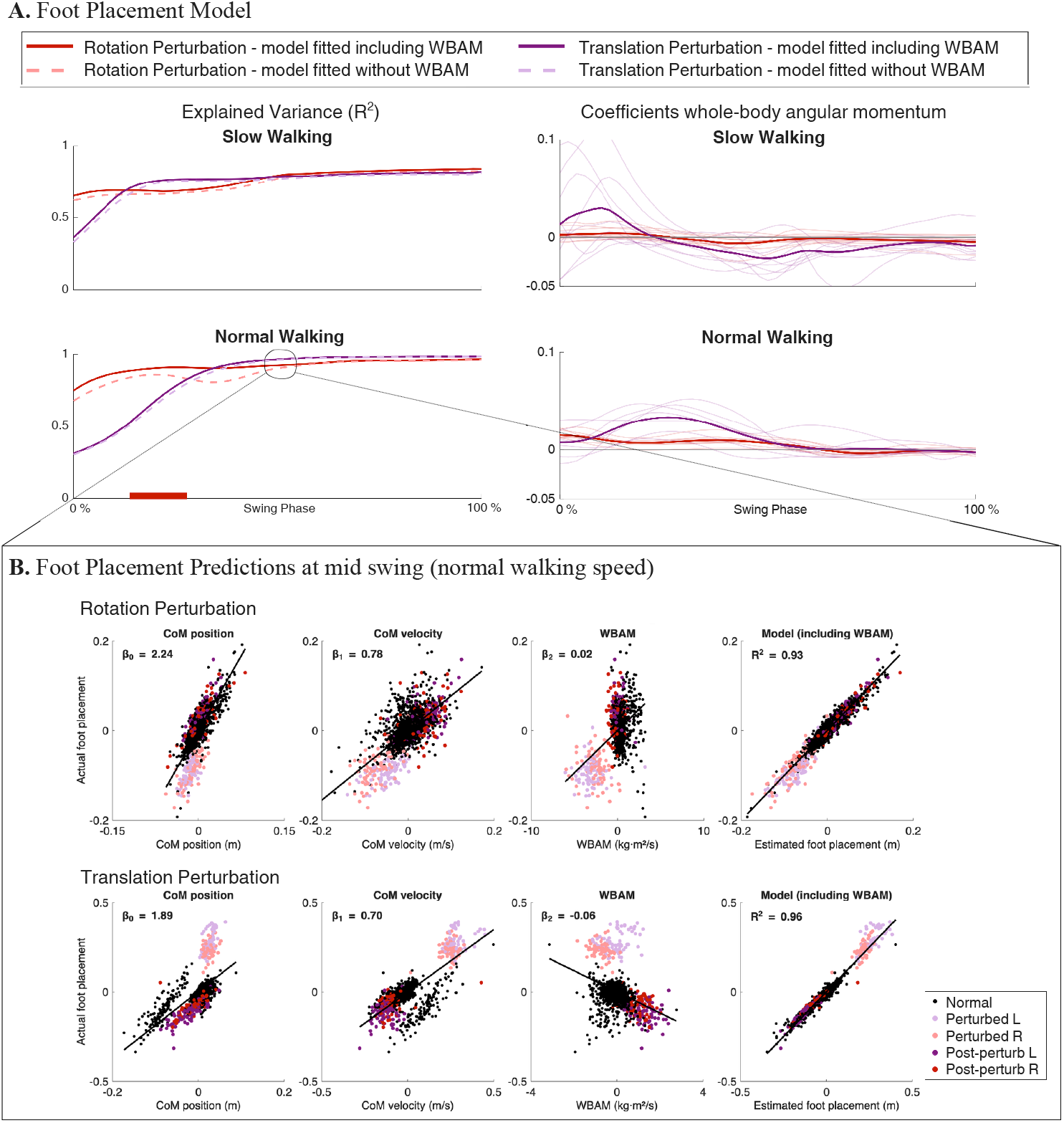
(A) Explained variance (R^2^; left figure) and regression coefficient whole-body angular momentum (WBAM, right figure) during the swing phase for mediolateral foot placement models for slow and normal walking speed. Thin transparent lines show the regression coefficient of whole-body angular momentum for individual participants. Thick solid line shows the group mean. Model performance is shown for translation (purple) and rotation (red) perturbations, for models excluding whole-body angular momentum as a predictor (dashed lines, foot placement model 1) and including whole-body angular momentum as a predictor (solid lines, foot placement model 2). The red box highlights the portion of the swing phase (∼10–20%) where including WBAM showed significantly higher explained variance for normal walking under rotation perturbations. (B) Relationship between mediolateral foot placement and each predictor variable (CoM position, CoM velocity, and whole-body angular momentum) at mid swing (50% of the swing phase), based on Model 2. The x-axis shows the predictor values at mid swing, and the y-axis shows the corresponding mediolateral foot placement at heel strike. The slope of each relationship reflects the regression coefficient for that predictor 50% of the swing phase. This panel provides a representative example of how each predictor is associated with foot placement. The arrow in panel A indicates the corresponding mid-swing time point.

Figure 5B illustrates the regression between mediolateral foot placement and each predictor at mid swing (50% of the swing phase), providing a representative example of how CoM position, CoM velocity, and whole-body angular momentum are associated with foot placement. Supplementary Figures S1–S12 present the regression relationships separately for perturbation type (translation vs. rotation), walking speed (normal vs. slow), and predictor (CoM position, CoM velocity, and whole-body angular momentum) at specific instances in the swing phase. Specifically, S1–S3 show translation perturbations during normal walking speed, S4–S6 show rotation perturbations during normal walking speed, S7–S9 show translation perturbations during slow walking speed, and S10–S12 show rotation perturbations during slow walking speed. Visual inspection of these figures shows that the association between CoM state and mediolateral foot placement becomes progressively stronger from early to late swing, with data aligning more clearly along a positive slope toward heel strike. In contrast, whole-body angular momentum demonstrates weaker and less consistent associations across swing, with only modest early-swing contributions following rotational perturbations.

## Discussion

We studied how foot placement is controlled following perturbations that affected either CoM linear momentum (translation perturbations) or whole-body angular momentum (rotation perturbations). We did so by comparing models which estimated deviations in foot placement based on CoM state alone with models that additionally included whole-body angular momentum. We found that foot placements models that included whole-body angular momentum as a predictor (figure 5A, solid red lines) only slightly increase the explained variance compared to models with only CoM kinematic information (Supplementary Figures S1–S12 and stable model coefficients in S14). This indicates that mediolateral foot placement is primarily coordinated with respect to CoM linear momentum, while whole-body angular momentum plays a limited role. This finding is consistent with models of stepping control that predict foot placement from the instantaneous CoM state (Bruijn & Van Dieën, 2018; Townsend, 1985; Wang & Srinivasan, 2014) and with experimental perturbation studies showing that mediolateral CoM velocity strongly predicts subsequent step location (Vlutters et al., 2016).

Contrary to our initial hypothesis, deviations in whole-body angular momentum explained only a small additional portion of the variance in foot placement, even in response to angular momentum perturbations. Participants appeared to restore whole-body angular momentum to near steady-state values during the perturbed stance phase, likely through modulations of mediolateral ground reaction forces. Such modulation of ground reaction forces during stance has previously been attributed to active muscle contributions, particularly from the hip abductors and ankle plantar flexors, which regulate whole-body angular momentum and body orientation (Herr & Popovic, 2008). Simulation studies have further demonstrated that stance-phase muscle activity can actively shape ground reaction forces and segmental angular momentum to stabilize gait (Neptune & McGowan, 2016). Such early whole-body angular momentum corrections are consistent with observations by Van Mierlo et al., (2022, 2023), who showed that whole-body angular momentum stabilization occurs very early after a disturbance and is driven largely by re-direction of the ground reaction force rather than by substantial center of pressure shifts. Their work also demonstrated that individuals prioritize restoring whole-body orientation before addressing translational CoM deviations.

The early restoration of whole-body angular momentum likely explains why including whole-body angular momentum improved foot placement predictions only during early swing in our data. Once whole-body angular momentum is sufficiently regulated during stance, the remaining control effort likely focuses on re-establishing CoM stability through appropriate foot placement. The relatively small contribution of whole-body angular momentum to foot placement modulation contrasts with findings from Leestma et al. (2023), who reported stronger whole-body angular momentum step-placement relationships, particularly in the frontal plane. These differences may reflect methodological factors: Leestma et al. (2023) perturbed participants later in stance and used integrated whole-body angular momentum over the full step, which masks in-stance corrections. In contrast, our perturbations were applied early, allowing participants to correct whole-body angular momentum before needing to adjust foot placement, thereby diminishing its apparent predictive value. Given this, perturbations delivered later in stance (e.g., mid-stance or terminal stance) may have prevented the rapid in-stance correction of whole-body angular momentum observed in our data, and whole-body angular momentum might then have explained a larger portion of foot placement variance. Leestma et al. (2023) used a treadmill translation as a perturbation which might result in a different perturbation magnitude and deviations in linear and angular momentum. Stronger perturbations may have elicited more pronounced responses, potentially impacting foot placement more significantly.

### Study limitations

Several methodological limitations should be considered when interpreting our results. First, perturbations were delivered only at heel strike, so it remains unclear how foot placement control and the contribution of whole-body angular momentum might differ for perturbations applied later in stance. Second, walking speed was fixed at two levels (slow and normal), which may limit the generalizability of our findings to faster or variable walking speeds. Third, the perturbation magnitude and direction were standardized, and the vertical height of the Bump’m system was fixed, which may have constrained the range of biomechanical responses. Fourth, although the perturbations were designed to selectively target either linear or angular momentum, complete isolation was not possible. The linear perturbations induced small but consistent changes in angular momentum, and conversely, the rotational perturbations induced small changes in linear momentum (Figure 3). This is caused by the fact that the body is in contact with the ground during the perturbation. For example, when approximating the body as an inverted pendulum, the translational perturbation will cause horizontal components of the ground reaction force through the pendulum dynamics, thereby inducing a perturbation of angular momentum. This mechanism can explain the small but consistent deviations in medio-lateral COM of mass position and velocity during the perturbation (Figure 3, S1-12). Finally, our sample consisted of healthy young adults, so caution is warranted when extrapolating these findings to populations with altered biomechanics, such as children, older adults, or individuals with obesity or neurological impairments (Kim et al., 2022). Future studies should systematically vary perturbation timing, magnitude, and speed, and examine diverse populations, to better understand the generalizability of these control strategies.

## Conclusions

Foot placement in the mediolateral direction is predominantly correlated with linear momentum, and only to a limited extent with whole-body angular momentum following both translation and rotation perturbations applied at heel strike. Our results indicate that rotational deviations are corrected rapidly within the stance phase, and that subsequent foot placement appears to be used to control linear momentum.

## Acknowledgements

Simone Berkelmans was supported by the Netherlands Organization for Scientific Research (NWO), grant OCENW.XS23.1.143. The authors would like to thank Lars van Rikxoord for his valuable assistance during this project. We also thank Richard Casius and Bert Clairbois for their support with the measurement (“bump’m”) system throughout the study.

