## Supplementary Material (figures and tables) for "DEVIATIONS IN WHOLE BODY ANGULAR MOMENTUM ARE LARGELY CORRECTED BEFORE FOOT PLACEMENT"

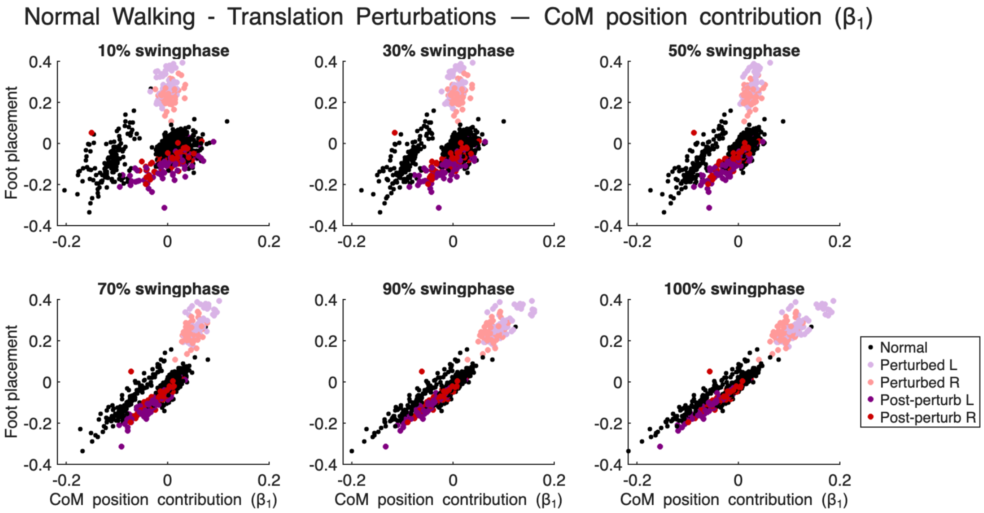


*Figure S1. Relationship between mediolateral foot placement and CoM position across swing phases during normal walking and translation perturbations. Each panel shows the relationship between CoM position and mediolateral foot placement at different percentages of the swing phase (10%, 30%, 55%, 70%, 90%, and 100%). The slope of the relationship reflects the regression coefficient ($\beta_1$). Black points represent steps during unperturbed normal walking. Colored points illustrate steps during perturbations (left or right) and the corresponding post-perturbation recovery steps.*

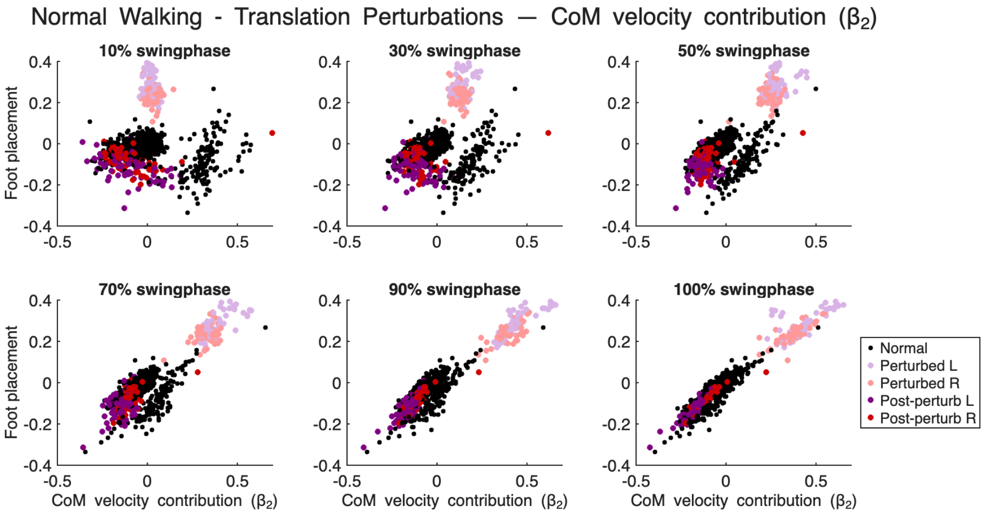


*Figure S2. Relationship between mediolateral foot placement and CoM velocity across swing phases during normal walking and translation perturbations. Each panel shows the relationship between CoM velocity and mediolateral foot placement at different percentages of the swing phase (10%, 30%, 55%, 70%, 90%, and 100%). The slope of the relationship reflects the regression coefficient ($\beta_2$). Black points represent steps during unperturbed normal walking. Colored points illustrate steps during perturbations (left or right) and the corresponding post-perturbation recovery steps.*

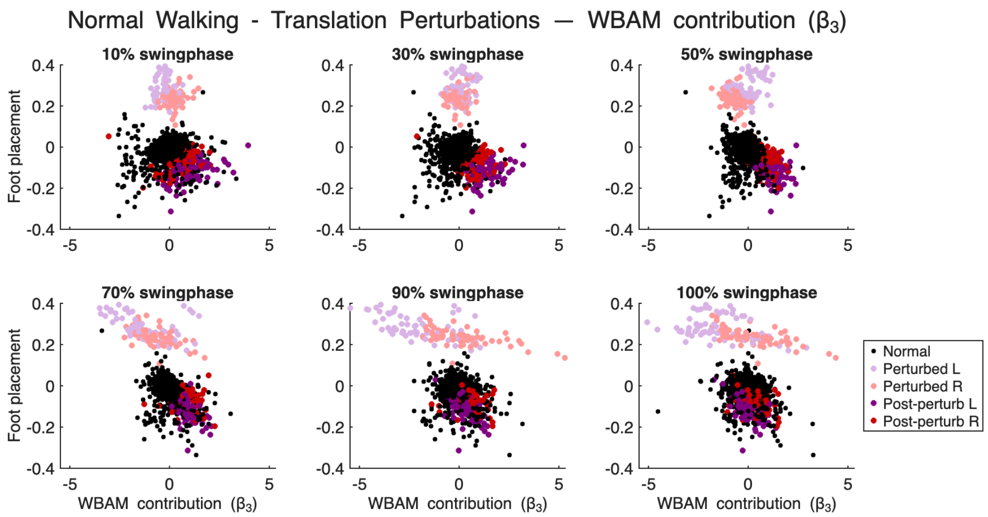


*Figure S3. Relationship between mediolateral foot placement and whole-body angular momentum (WBAM) across swing phases during normal walking and translation perturbations. Each panel shows the relationship between WBAM and mediolateral foot placement at different percentages of the swing phase (10%, 30%, 55%, 70%, 90%, and 100%). The slope of the relationship reflects the regression coefficient ($\beta_3$). Black points represent steps during unperturbed normal walking. Colored points illustrate steps during perturbations (left or right) and the corresponding post-perturbation recovery steps.*

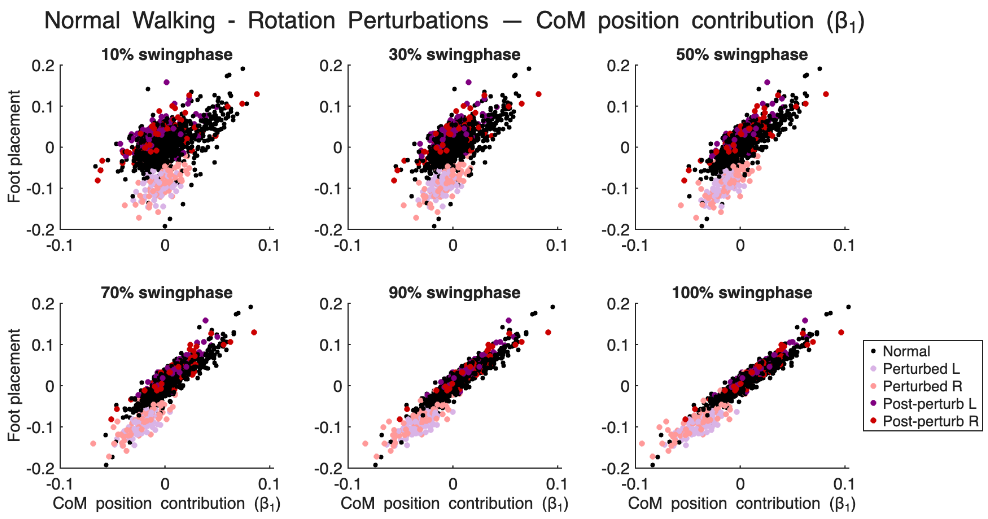


*Figure S4. Relationship between mediolateral foot placement and CoM position across swing phases during normal walking and rotation perturbations. Each panel shows the relationship between CoM position and mediolateral foot placement at different percentages of the swing phase (10%, 30%, 55%, 70%, 90%, and 100%). The slope of the relationship reflects the regression coefficient ($\beta_1$). Black points represent steps during unperturbed normal walking. Colored points illustrate steps during perturbations (left or right) and the corresponding post-perturbation recovery steps.*

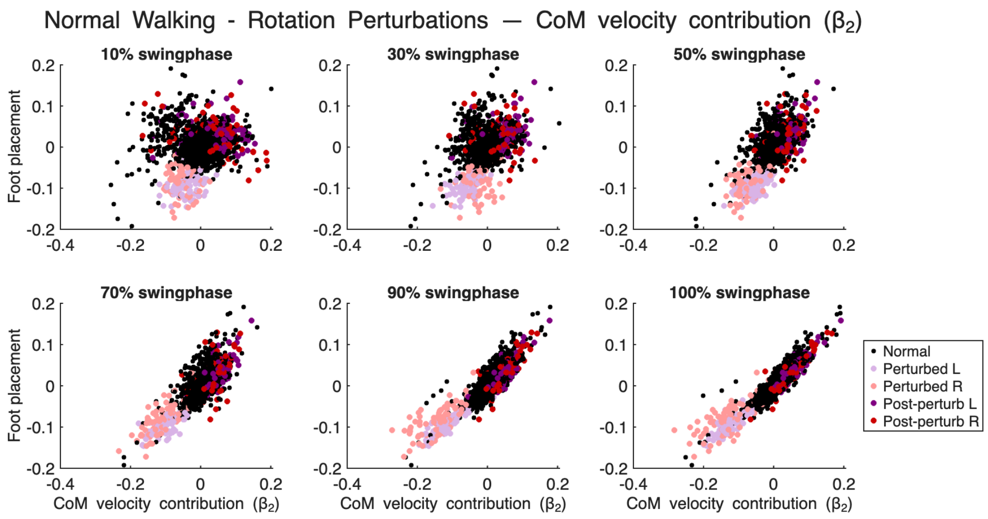


*Figure S5. Relationship between mediolateral foot placement and CoM velocity across swing phases during normal walking and rotation perturbations. Each panel shows the relationship between CoM velocity and mediolateral foot placement at different percentages of the swing phase (10%, 30%, 55%, 70%, 90%, and 100%). The slope of the relationship reflects the regression coefficient ($\beta_2$). Black points represent steps during unperturbed normal walking. Colored points illustrate steps during perturbations (left or right) and the corresponding post-perturbation recovery steps.*

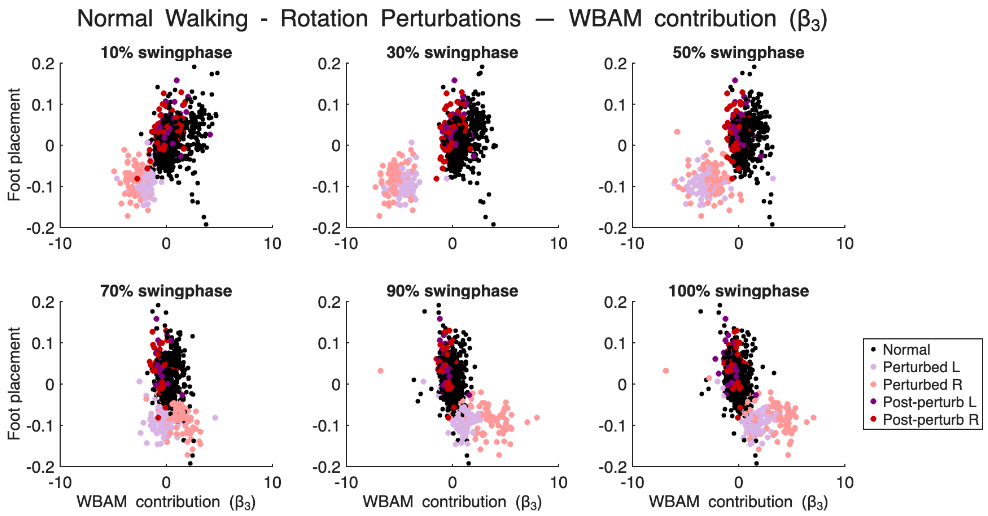


*Figure S6. Relationship between mediolateral foot placement and whole-body angular momentum (WBAM) across swing phases during normal walking and rotation perturbations. Each panel shows the relationship between WBAM and mediolateral foot placement at different percentages of the swing phase (10%, 30%, 55%, 70%, 90%, and 100%). The slope of the relationship reflects the regression coefficient ($\beta_3$). Black points represent steps during unperturbed normal walking. Colored points illustrate steps during perturbations (left or right) and the corresponding post-perturbation recovery steps.*

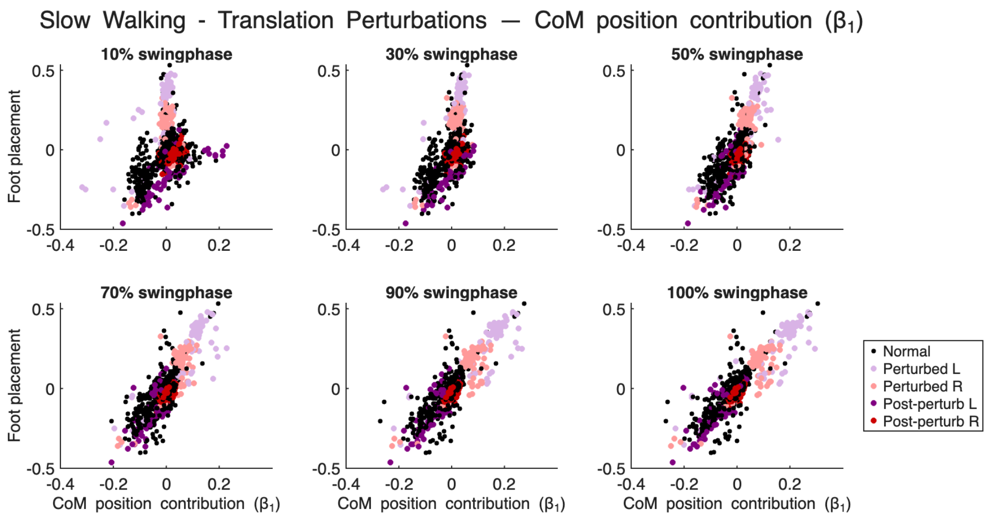


*Figure S7. Relationship between mediolateral foot placement and CoM position across swing phases during slow walking and translation perturbations. Each panel shows the relationship between CoM position and mediolateral foot placement at different percentages of the swing phase (10%, 30%, 55%, 70%, 90%, and 100%). The slope of the relationship reflects the regression coefficient ($\beta_1$). Black points represent steps during unperturbed slow walking. Colored points illustrate steps during perturbations (left or right) and the corresponding post-perturbation recovery steps.*

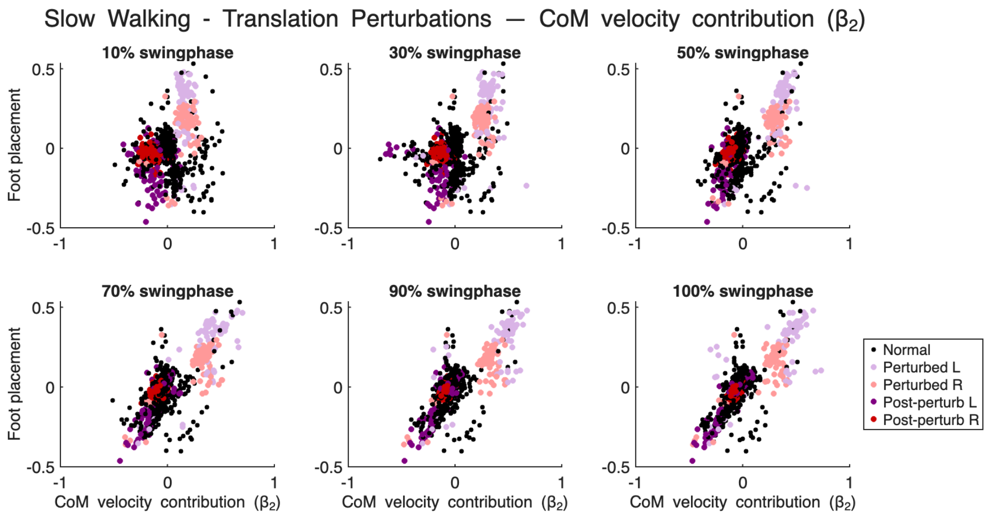


*Figure S8. Relationship between mediolateral foot placement and CoM velocity across swing phases during slow walking and translation perturbations. Each panel shows the relationship between CoM velocity and mediolateral foot placement at different percentages of the swing phase (10%, 30%, 55%, 70%, 90%, and 100%). The slope of the relationship reflects the regression coefficient ($\beta_2$). Black points represent steps during unperturbed slow walking. Colored points illustrate steps during perturbations (left or right) and the corresponding post-perturbation recovery steps.*

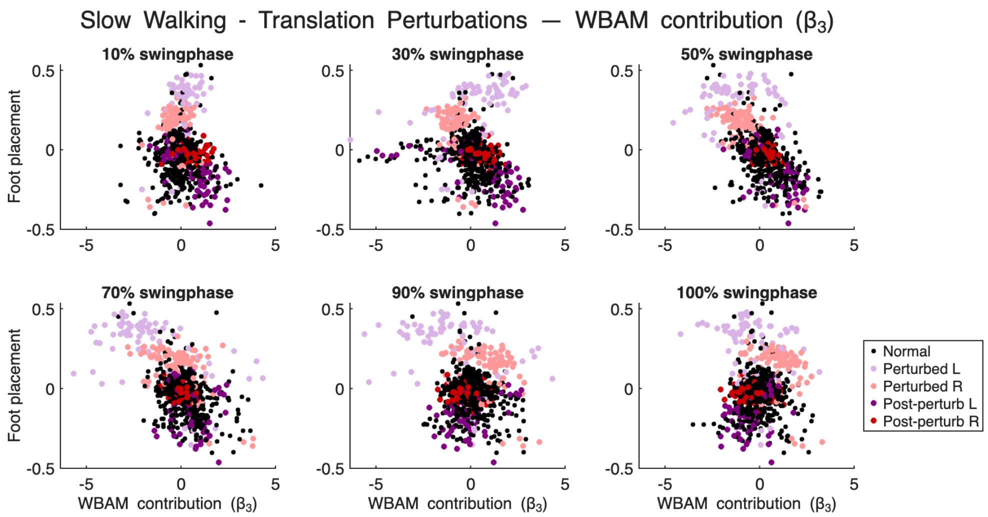


*Figure S9. Relationship between mediolateral foot placement and whole-body angular momentum (WBAM) across swing phases during slow walking and translation perturbations. Each panel shows the relationship between WBAM and mediolateral foot placement at different percentages of the swing phase (10%, 30%, 55%, 70%, 90%, and 100%). The slope of the relationship reflects the regression coefficient ($\beta_3$). Black points represent steps during unperturbed slow walking. Colored points illustrate steps during perturbations (left or right) and the corresponding post-perturbation recovery steps.*

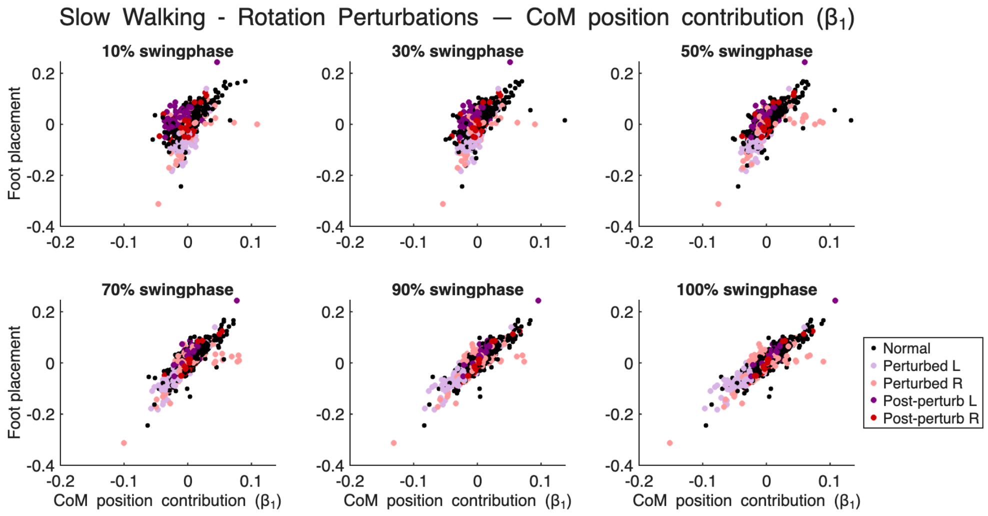


*Figure S10. Relationship between mediolateral foot placement and CoM position across swing phases during slow walking and rotation perturbations. Each panel shows the relationship between CoM position and mediolateral foot placement at different percentages of the swing phase (10%, 30%, 55%, 70%, 90%, and 100%). The slope of the relationship reflects the regression coefficient ($\beta_1$). Black points represent steps during unperturbed slow walking. Colored points illustrate steps during perturbations (left or right) and the corresponding post-perturbation recovery steps.*

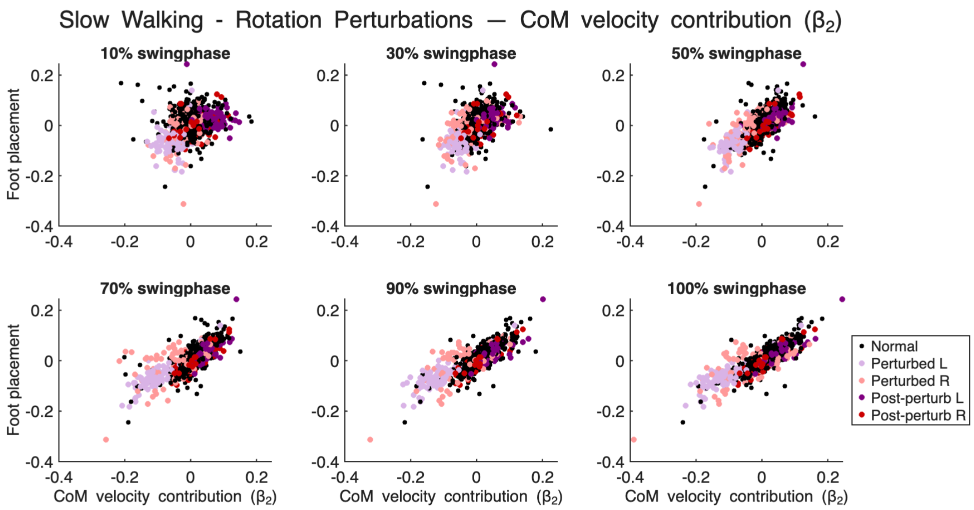


*Figure S11. Relationship between mediolateral foot placement and CoM velocity across swing phases during slow walking and rotation perturbations. Each panel shows the relationship between CoM velocity and mediolateral foot placement at different percentages of the swing phase (10%, 30%, 55%, 70%, 90%, and 100%). The slope of the relationship reflects the regression coefficient ($\beta_2$). Black points represent steps during unperturbed slow walking. Colored points illustrate steps during perturbations (left or right) and the corresponding post-perturbation recovery steps.*

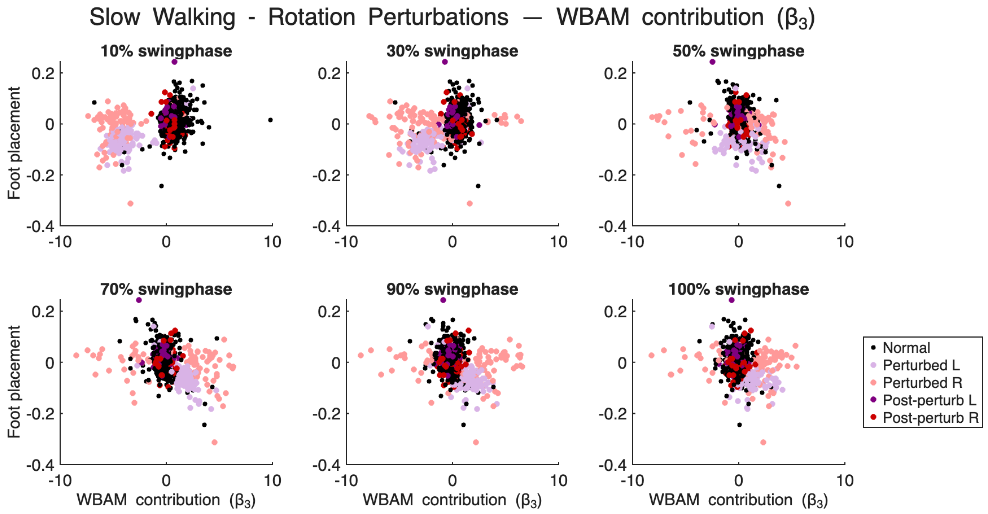


*Figure S12. Relationship between mediolateral foot placement and whole-body angular momentum (WBAM) across swing phases during slow walking and rotation perturbations. Each panel shows the relationship between WBAM and mediolateral foot placement at different percentages of the swing phase (10%, 30%, 55%, 70%, 90%, and 100%). The slope of the relationship reflects the regression coefficient ($\beta_3$). Black points represent steps during unperturbed slow walking. Colored points illustrate steps during perturbations (left or right) and the corresponding post-perturbation recovery steps.*

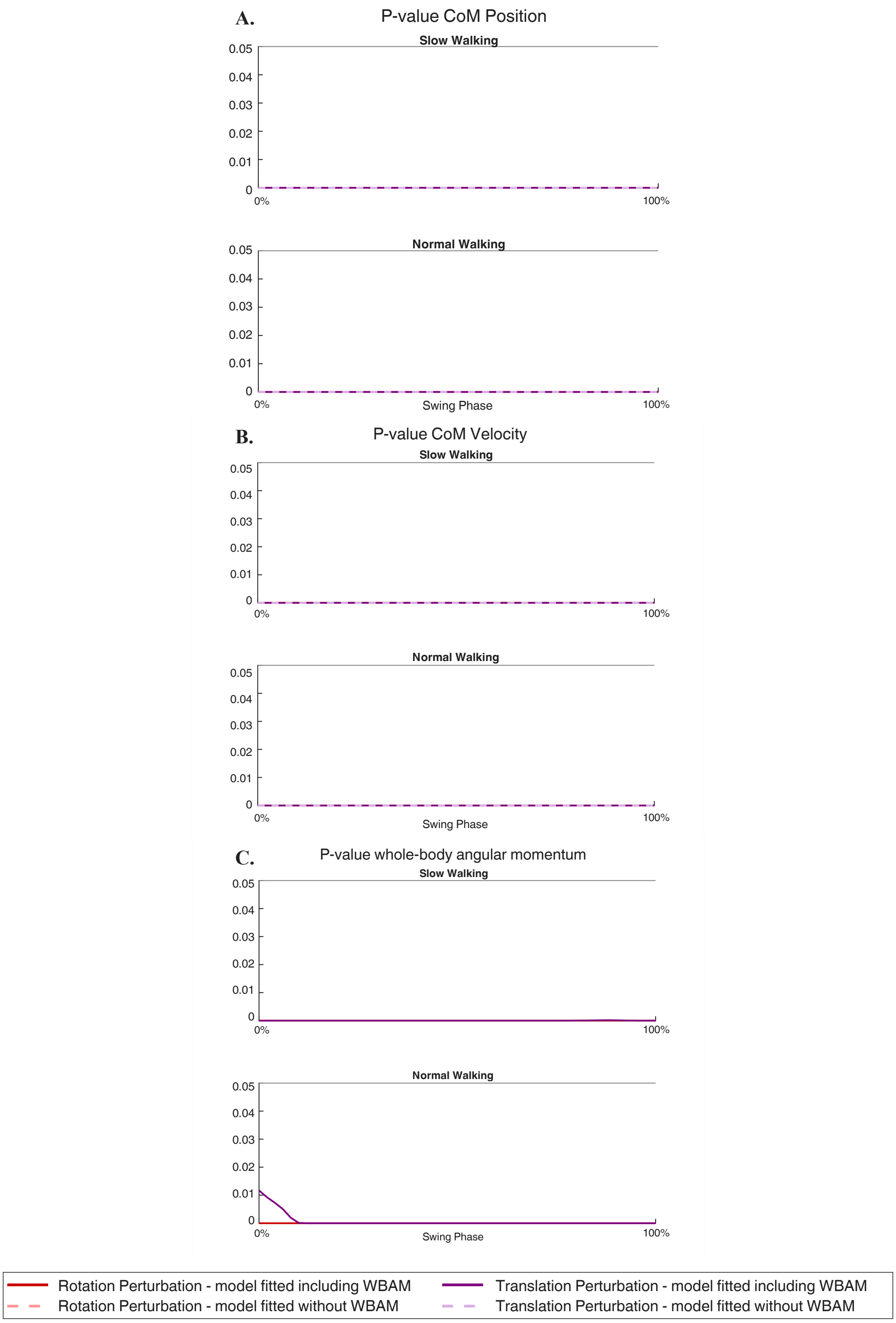


*Figure S13. P-values of regression coefficients across the swing phase for CoM position (A), CoM velocity (B), and whole-body angular momentum (C) during slow and normal walking.*

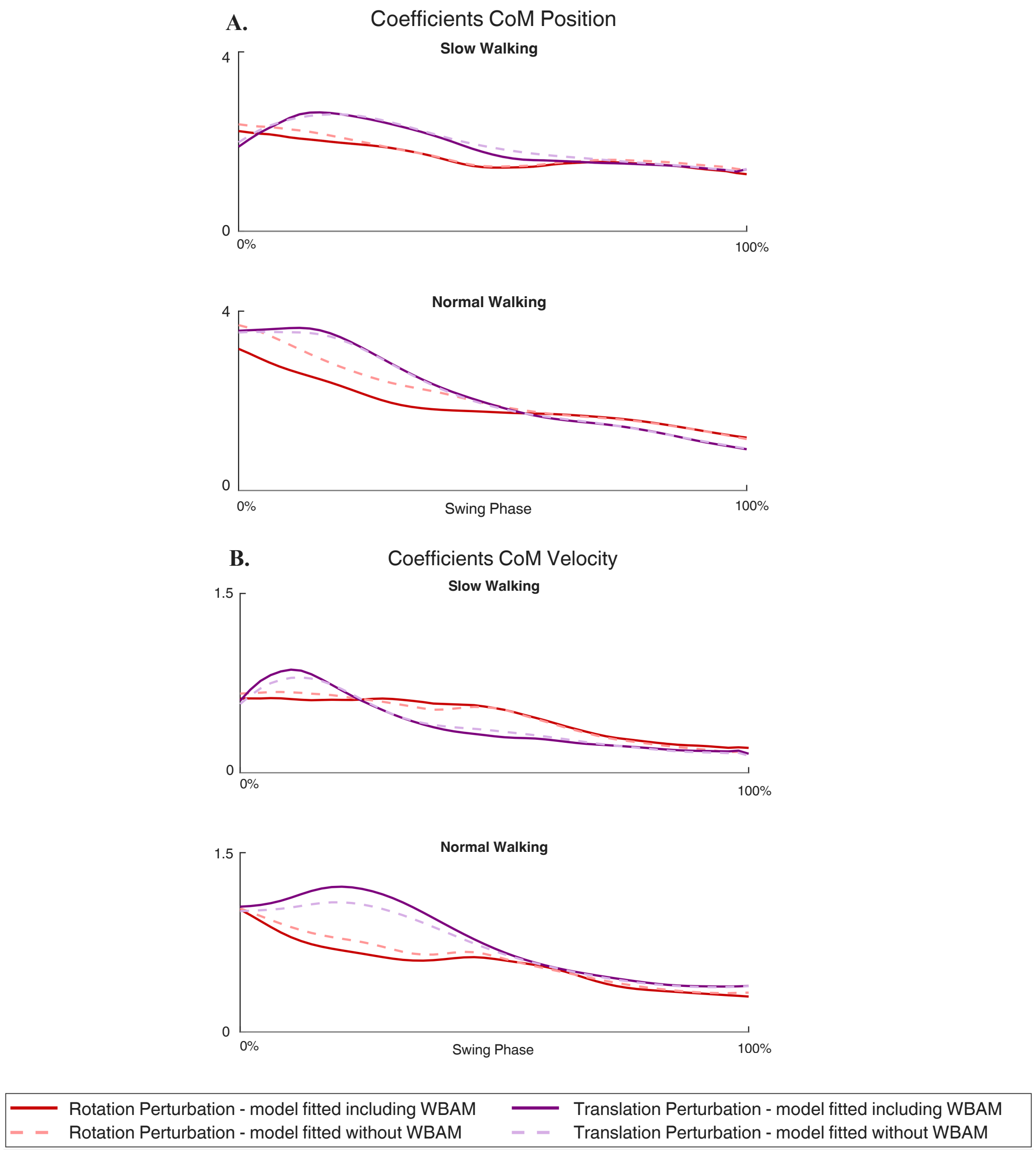


*Figure S14. Coefficient of (A) CoM Position and (B) CoM Velocity of mediolateral foot placement models during the swing phase for slow and normal walking speed. Panels show the estimated regression coefficients $\beta_1$ (CoM position) and $\beta_2$ (CoM velocity) derived from two foot-placement models. Model 1 included only CoM position and velocity (red and purple dashed lines), and Model 2 additionally incorporated whole-body angular momentum (red and purple solid lines).*

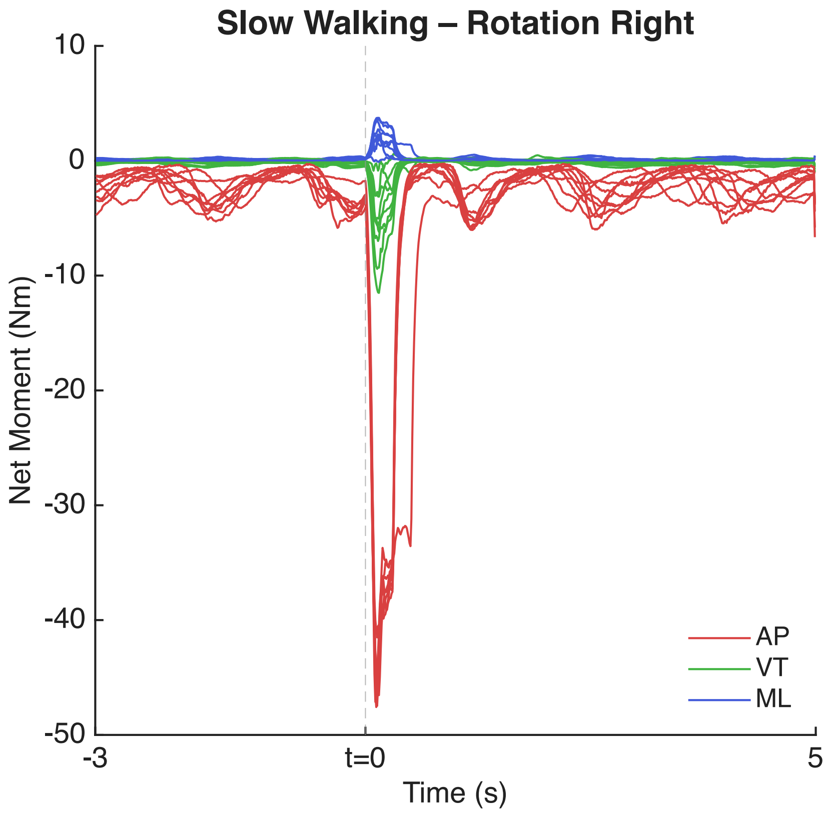


*Figure S15*. M*oments about the CoM during rotation perturbations at slow walking speed, plotted with perturbation onset at t = 0, including 3 seconds prior and 5 seconds following the perturbation. Each line represents a participant. AP: anterior-posterior, ML: mediolateral, and VT: vertical.*

**Supplementary Material - Tables**

*Table S1 - Placement of (cluster) markers on the segments/bony landmarks*

**Torso Markers**

| | | |
|---|---|---|
| C7 | 7th Cervical Vertebrae | Spinous process on the 7th cervical vertebrae |
| HARNESS (C7_2) | On the safety harness | Marker on the safety harness, around T5 |

**Arm Markers**

| | | |
|---|---|---|
| R/LSHO | Right/Left Shoulder | On the acromio-clavicular joint |
| R/LUA 1-3 | Right/Left Upper Arm (cluster) | The upper arm between shoulder and elbow markers |
| R/LELB_lat | Right/Left Lateral Epicondyle | |
| R/LLA 1-3 | Right/Left Lower Arm (cluster) | The lower arm between elbow and wrist markers |
| R/LWRA | Styloid Process (Radius) | |
| R/LWRB | Styloid Process (Ulna) | |

**Pelvis Markers**

| | | |
|---|---|---|
| R/LASIS | Right/Left ASIS | Right/Left anterior superior iliac spine |
| R/LPSIS | Right/Left PSIS | Right/Left posterior superior iliac spine |

**Leg Markers**

| | | |
|---|---|---|
| R/LFEM 1-3 | Right/Left Thigh/Femoris (cluster) | Placed on the lower lateral surface of the thigh |
| R/LKNEE_med | Right/Left Medial Epicondyle | |
| R/LKNEE_lat | Right/Left Lateral Epicondyle | |
| R/LTIB 1-3 | Right/Left Shank (cluster) | Placed on the lateral side of the shank |
| R/LANK_med | Right/Left Malleolus Medialis | |
| R/LANK_lat | Right/Left Malleolus Lateralis | |
| R/LCAL | Right/Left Calcaneus | |
| R/LMTP5 | Right/Left foot tuberositas ossis metatarsi | Tuberositas ossis metatarsi |

**Bump’m Markers**

| | | |
|---|---|---|
| RPEL_bump | Bump’m String Pelvis | Placed where the string is attached to the pelvis |
| R_BUMP | Right Bump’m String | Placed on the string where the force sensor is placed |
| LSHO_bump | Bump’m String Shoulder | Placed where the string is attached to the shoulder |
| L_BUMP | Left Bump’m String | Placed on the string where the force sensor is placed |
